# Second language learning in older adults modulates Stroop task performance and brain activation

**DOI:** 10.1101/2024.03.07.583963

**Authors:** DH Schultz, A Gansemer, K Allgood, M Gentz, L Secilmis, Z Deldar, CR Savage, L Ghazi Saidi

**Affiliations:** Department of Psychology, University of Nebraska-Lincoln; Center for Brain, Biology, and Behavior, University of Nebraska-Lincoln; Department of Communication Disorders, College of Education, University of Nebraska at Kearney (UNK); Department of Psychology, McGill University, Montreal, Canada

**Keywords:** Language Learning, Cognitive Effects, Stroop task, Older Adults, Aging, Cognitive Reserve, fMRI, Neuroimaging, Neurocognition

## Abstract

Numerous studies have highlighted cognitive benefits in lifelong bilinguals during aging, manifesting as superior performance on cognitive tasks compared to monolingual counterparts. Yet, the cognitive impacts of acquiring a new language in older adulthood remain unexplored. In this study, we assessed both behavioral and fMRI responses during a Stroop task in older adults, pre- and post language-learning intervention.

A group of 41 participants (age:60-80) from a predominantly monolingual environment underwent a four-month online language course, selecting a new language of their preference. This intervention mandated engagement for 90 minutes a day, five days a week. Daily tracking was employed to monitor progress and retention. All participants completed a color-word Stroop task inside the scanner before and after the language instruction period.

We found that performance on the Stroop task, as evidenced by accuracy and reaction time, improved following the language learning intervention. With the neuroimaging data, we observed significant differences in activity between congruent and incongruent trials in key regions in the prefrontal and parietal cortex. These results are consistent with previous reports using the Stroop paradigm. We also found that the amount of time participants spent with the language learning program was related to differential activity in these brain areas. Specifically, we found that people who spent more time with the language learning program showed a greater increase in differential activity between congruent and incongruent trials after the intervention relative to before.

Future research is needed to determine the optimal parameters for language learning as an effective cognitive intervention for aging populations. We propose that with sufficient engagement, language learning can enhance specific domains of cognition such as the executive functions. These results extend the understanding of cognitive reserve and its augmentation through targeted interventions, setting a foundation for future investigations.

## Introduction

In recent years, the prevention of cognitive decline in normal or pathological aging has emerged as a paramount focus of many neuropsychological studies (Di Nuovo et al., 2020). This is particularly important given the increasing aging population (Kanasi et al., 2016) and the prevalence of age-related cognitive disorders (Lopez & Kuller, 2019). The role of nonpharmaceutical interventions in mitigating the progression of cognitive decline is crucial.

Unlike pharmacological treatments, which often target specific symptoms or stages of cognitive diseases, nonpharmaceutical interventions offer a holistic approach with the potential for broader applicability and fewer side effects (Dyer et al., 2018). Non-pharmaceutical interventions which encompass a variety of options from lifestyle modifications to cognitive training, and dietary adjustments are considered proactive strategies to preserve cognitive function (Klimova et al., 2017) and strengthen cognitive reserve (Bott et al., 2019).

Cognitive reserve, a fundamental concept in healthy aging, refers to the brain’s resilience to neuropathological damage (Cammisuli et al., 2022; Scarmeas & Stern, 2003). Emerging evidence suggests that higher cognitive reserve is associated with greater ability to compensate for age-related brain changes and pathology, thereby mitigating the manifestation of clinical symptoms (M. Tucker & Stern, 2011). This resilience is believed to stem from a variety of life-long experiences, including education, occupational complexity, and engagement in cognitively stimulating activities such as music, games, reading and so forth (Liberati et al., 2012).

Among the factors contributing to cognitive reserve, bilingualism has emerged as a significant area of interest in the past decade (Bialystok, 2021). There is behavioral and neuroimaging evidence that lifelong bilingualism enhances cognitive reserve (Craik et al., 2010; Schweizer et al., 2012). Bilingual individuals engage in constant cognitive exercises, such as switching between languages and inhibiting one language while using another (Green & Abutalebi, 2013). This cognitive exercise over a lifetime is believed to strengthen neural networks and enhance cognitive flexibility (Barbu et al., 2018). Further, there is evidence that the cognitive demand associated with managing two languages contributes to the development of higher cognitive reserve (Bialystok et al., 2007) and therefore delaying the onset of age-related cognitive decline and neurodegenerative diseases (Bialystok et al., 2016).

Many researchers agree on the underlying reasons why bilingualism might contribute to enhanced cognitive reserve. Engaging in bilingualism requires the navigation of complex linguistic structures, managing two linguistic systems simultaneously, requiring inhibitory control and therefore continuously engaging executive control functions, such as task switching and inhibition (Kroll & Bialystok, 2013). These cognitive demands are hypothesized to lead to structural and functional brain changes, including in areas involved in executive control and language processing (García-Pentón et al., 2014). Additionally, the constant management of dual language systems is believed to enhance neural plasticity and efficiency. This may provide a neurological buffer against age-related cognitive decline (Del Maschio et al., 2018).

Despite the body of evidence for cognitive advantages of lifelong bilingualism in aging, research on the cognitive effects of second language (L2) acquisition in older adults is still in its infancy. A few studies have reported that L2 learning in older adults can lead to improvements in global cognition (Antoniou et al., 2013) and specific cognitive tasks such as memory, attention, and executive functioning (Meltzer et al., 2023a), potentially offering a protective mechanism against cognitive decline. This enhancement is often attributed to the cognitive stimulation and neural plasticity induced by the complexities of acquiring a new language (P. Li et al., 2014), which in turn may boost cognitive reserves (Antoniou et al., 2013).

The cognitive demands of second language (L2) acquisition is well established by theoretical models such as the dynamic model (Abutalebi & Green, 2007), and experimental research (Fukuta & Yamashita, 2015). Learning a new language entails extensive engagement of various cognitive systems, particularly those involved in memory, attention, and executive functioning (Ghazi Saidi et al., 2013; Ghazi-Saidi & Ansaldo, 2017). This engagement is hypothesized to not only activate and strengthen these cognitive areas but also potentially induce neuroplastic changes within the brain (Ghazi-Saidi et al., submitted).

The Stroop task is an excellent choice for measuring the cognitive effects of language learning for several reasons. First, the Stroop task is designed to assess executive function, particularly cognitive control and inhibition (Egner & Hirsch, 2005; Guarino et al., 2020). Learning a new language requires similar cognitive skills, such as controlling attention between different language systems and inhibiting one language system while using another (Green & Abutalebi, 2013; Hirosh & Degani, 2018; Kroll et al., 2008). The Stroop task is also known to activate several brain regions associated with language processing, attention, and executive function (Heidlmayr et al., 2020; Ye & Zhou, 2009). Therefore, learning a new language may improve performance on the Stroop task. Second, given that Stroop task involves processing words and colors, it inherently taps into language processing areas of the brain (Hertrich et al., 2021).

Thus, the Stroop task provides an objective measure to assess a specific cognitive domain at baseline and after the intervention. Any significant change in performance on the Stroop task from pre to post intervention can be attributed to cognitive changes potentially induced by the language learning process.

## Methods

### Design and Procedure

In this study, we used a pre-post intervention design with functional magnetic resonance imaging (fMRI) to explore the neural effects of language learning in older adults. The study was initiated by conducting baseline fMRI scans for all participants to establish a neurological benchmark. Following this, the participants underwent an online language learning program.

Participants were monitored daily for their performance and retention. After completion of the intervention, a second set of fMRI scans was conducted to identify any neural changes attributable to the intervention. This pre-post design allows for the assessment of neural adaptations in response to new language acquisition in older adults. The fMRI data were analyzed using standard neuroimaging techniques to observe changes in brain activity, with a particular focus on regions associated with language processing and cognitive control.

This study was approved by the UNL Institutional Review Board (IRB), Research Compliance Services, Office of Research & Economic Development, University of Nebraska-Lincoln. The University of Nebraska at Kearney IRB acknowledged and honored the site agreement to cede IRB review to the University of Nebraska-Lincoln IRB regarding this study, under the SMART IRB Master Common Reciprocal Institutional Review Board Authorization Agreement. Per this agreement the UNL IRB serves as the reviewing IRB and the UNK IRB as the relying IRB.

### Participants

Our participant cohort comprised 41 healthy, monolingual individuals, aged between 60 to 80 years (M= 66.63, SD=4.7, min=60; max=77), all residing in a predominantly monolingual environment in Nebraska. All participants had at least 14 years of education (M=17.5, SD=2.94). All participants were white caucasian, mirroring the demographics of the rural Nebraskan population. All ethnic minorities were excluded based on proficiency in two or more languages.

### Inclusion and Exclusion Criteria

Inclusion criteria consisted of being monolingual English-speaking adults, aged 60 - 80 years of any gender, race or ethnicity, with an electronic device and access to the internet, with normal or corrected vision and hearing, no memory or learning problems, no diagnosed depression or neurological disorders. In addition, we only included right-handed individuals to control for left-hemisphere dominance. All participants were fully vaccinated for Covid-19 given that data were collected shortly after Covid-19 restrictions were lifted to avoid putting older adults at additional risk of exposure to Covid-19. Participants who were claustrophobic or did not have the ability to report to the Center for Brain, Biology and Behavior in Lincoln, NE, or did not pass the MRI compatibility screener were excluded. All participants had computer skills to enroll in the intervention program and access to the internet and a device. All participants were screened using Montreal Cognitive Assessment (MoCA) and participants who did not score above the cut-off of 26 were excluded.

The Language Experience & Proficiency Questionnaire (LEAP-Q) (Kaushanskaya et al., 2020; Marian et al., 2007) was used to screen for second language knowledge, use and exposure. LEAP-Q is a self-report toolbox to collect data on language knowledge, use and exposure for all languages spoken by an individual. Scores range 0-10, and scores above 7 are taken as a measure of bilingual proficiency; a score of 0 indicates monolingualism. Only participants with a score of 0 or 1 (minimal knowledge and at vocabulary level) were included.

### Intervention

Participants engaged in an online language learning program, Rosetta Stone (https://www.rosettastone.com/), in which they selected a language of their choice to learn. Rosetta Stone is a comprehensive computer-assisted language learning software developed by Rosetta Stone Ltd. This software employs an immersive method, inspired by the naturalistic way individuals learn their first language (Work, 2014; Zheng, 2024). The approach is characterized by the absence of translations or explicit grammar instructions, instead relying on visual, auditory, and textual cues in the target language to convey meaning and foster language comprehension and production. This methodology aligns with the communicative approach to language teaching, which emphasizes the importance of interaction and using the language for real-life communication purposes. We used the educational version of Rosetta Stone which provided us with the possibility of monitoring the adherence to the program with detailed information about the number of minutes engaged in the program, a detailed statistics about the activities in which the participant engaged, the number of times each activity was repeated and scores on tests at the end of each lesson and each level. Among the 26 different options available, participants selected Spanish, French, German, Italian, and Japanese, in order of popularity.

The intervention spanned a duration of four months. Participants were instructed to engage in language learning five days per week, at a dose of 90 minutes per day. The online platform provided flexibility for participants to learn at their own pace and in a familiar environment, potentially enhancing adherence to the program. To ensure a consistent and effective learning experience, we offered optional monthly zoom meetings to participants. These meetings provided them with an opportunity to meet other participants, ask questions and share their experiences. Adherence to the intervention protocol was closely monitored. As administrators, we had access to log-in and time and type of language learning activities. We collected data on daily activities, and test results. At the end of each lesson, there was a test. Participants could proceed with the lesson only if they passed the test with 80% accuracy or above. This data was then used to assess the fidelity of the intervention and its potential impact on the cognitive abilities of the participants.

### Stroop Task

The color-word Stroop task (Scarpina & Tagini, 2017; Stroop, 1935) is a well-established neuropsychological test designed to evaluate cognitive control and executive function (Bari & Robbins, 2013; Kane & Engle, 2003). During the Stroop task, participants are presented with words denoting colors that are printed in congruent or incongruent colors (e.g., the word "red" printed in blue). They were required to identify the color of the “ink”, not the word itself, which requires the inhibition of an automatic reading response. Specifically, the congruent trials included the names of colors that appeared in the same color that they read. The incongruent trials included the names of colors that appeared in a different color that they read. The participants were instructed to always pick the color of the word on the monitor (i.e. the “ink”), and ignore what the word “read”. The neural condition consisted of the words “when,” “so” and “for” which were presented in different colors on the monitor. Participants were instructed to respond with a right index finger key press for when the color of the word was yellow, a right middle finger key press when the color of the word was red, and a right ring finger key press when the color of the word was green. The task was displayed to participants via a screen visible through a mirror mounted on the head coil.

The Stroop task paradigm was previously used in a number of studies (Kane & Engle, 2003; MacLeod, 1991). Our custom version of the Stroop task consisted of 108 trials, divided into three conditions with each condition comprising 36 trials. Stimuli were presented with E-prime version 2.0 software (Schneider et al., 2002). Trials were presented in a pseudorandom order to control for sequence effects: 1) Neutral Condition: Departing from the traditional non-word letter strings, our task incorporated common English words such as "When," "So," and "Like." These words, which do not inherently relate to colors, were presented in yellow, red, or green. This adaptation aimed to control the impact of familiarity and semantic content on the participants’ response times and accuracy. 2) Congruent Trials: For these trials, color words such as "Yellow," "Red," and "Green" were displayed in their respective colors. The semantic and visual congruence would expedite response times, leveraging the alignment between the word meaning and its visual presentation. 3) Incongruent Trials: These trials featured color words in contrasting colors, thereby inducing the Stroop effect. Each color word was repeated 12 times in each of the three incongruent colors, resulting in a total of 36 trials for each condition.

Each trial commenced with blank screen for 1,000 milliseconds (ms) to eliminate any afterimages or lingering visual distractions. Subsequently, a fixation signal was displayed for 200 ms to centralize the participant’s gaze and stabilize their visual field in preparation for the stimulus. The target letter string was then presented at the center of the screen for 1,500 ms, allowing a uniform duration for participant responses. Post-response, the inter-trial interval (ITI) varied randomly between 3,000 to 7,000 ms prior to the onset of the next stimulus. This variable ITI was implemented to prevent participants from anticipating the timing of stimuli, ensuring that responses were spontaneous. Response times and accuracy were recorded by the E-prime software. These metrics are integral for analyzing the Stroop effect’s magnitude (Kane & Engle, 2003; MacLeod, 1991).

Prior to entering the MRI environment, we explained the task to the participants. Participants also practiced a short version of the same task outside of the MRI environment. The only differences from the MRI Stroop task was that participants were seated at a desk and used a keyboard to respond.

### MRI Acquisition

MRI data were collected using a 3T Siemens Skyra scanner with a 32-channel head coil at the Center for Brain, Biology and Behavior at the University of Nebraska-Lincoln. A T1-weighted high resolution anatomical scan (TR = 2.2 s, TE = 3.37 ms, flip angle = 7°, FOV = 256 mm/100% phase, 192 slices, slice thickness = 1 mm) was collected for precise alignment of functional data. We also collected multiband echo-planar data (TR = 1 s, TE= 29.8 ms, flip angle = 60°, FOV = 210 mm/100% phase, slice thickness = 2.5 mm, 51 interleaved slices, multiband acceleration factor = 3) during in-scanner task performance (one run of 490 s prior to language learning and one run of 490 s following the language learning intervention).

### fMRI Preprocessing

Neuroimaging data were reconstructed using dcm2niix (Li et al., 2016). Following reconstruction, data preprocessing was completed using Analysis of Functional Neuroimages (AFNI) software (Cox, 1996). Code for the neuroimaging data processing and analysis is available at: https://github.com/dhschultz29/L2_learning_in_older_adults. Preprocessing consisted of despiking, slice time correction, non-linear transformation of anatomical data to MNI152_2009 template space, alignment of functional data to the transformed anatomical data, volume registration in which each volume is registered to the volume with the minimum outlier fraction, spatial smoothing using a 4 mm full-width at half maximum Gaussian filter, scaling the mean of each voxel’s time course to 100, and using a general linear model with three task regressors (correct congruent trials, correct incongruent trials, and correct neutral trials), twelve motion estimates (3 planes, 3 rotations, and their derivatives) from volume registration as regressors of no interest, and up to a sixth-order polynomial to model baseline and drift, all completed with the afni_proc.py function. We modeled hemodynamic response functions using the “BLOCK” basis function at the onset of each correct trial of each condition. We set the duration of the function to 1 sec. Consecutive pairs of volumes where the Euclidean norm of the motion derivatives exceeded 0.4 were “scrubbed” and eliminated from analysis (Power et al., 2012) along with the first 4 TRs of the run. The mean beta weight for each condition was extracted for each participant, voxel, and time point for subsequent statistical analysis.

### Data analysis

#### Behavioral Data Analysis

We calculated mean accuracy and reaction time data for congruent and incongruent trials for each participant. We ran an ANOVA with time (pre and post-language learning) and condition (congruent and incongruent) as repeated measures in JASP (ver. 0.16.2) (Love et al., 2019). The Holm-Bonferroni method was used for post hoc tests.

#### fMRI Data Analysis

We compared activation estimates for congruent and incongruent trials prior to the language learning intervention and following the language learning intervention using the 3dMVM function in AFNI (Chen et al., 2014). Participant was considered a random factor, condition and time point were modeled as within participant factors. We set a voxel-wise p-value threshold of 0.0001 for the main effects of condition (congruent/incongruent), time (pre-intervention, post-intervention), and the interaction of condition and time. We used a cluster-based approach to account for multiple comparisons for each of the main effects and the interaction (Forman et al., 1995). We estimated the smoothness of the residual time series and calculated the mean spatial autocorrelation parameters (ACF) across participants at pre-intervention (mean ACF = 0.7946, 2.1355, 4.8919) (Cox et al., 2017) and ten thousand random maps with these smoothness parameters were generated and thresholded at a voxelwise p < 0.001. The largest surviving cluster for each of these simulations was recorded and used to estimate the probability of a false positive. Based on these estimates we applied a cluster threshold to our data at a voxel-wise p-value of 0.001 and a minimum cluster size of seven voxels sharing either a face or an edge (NN = 2). This results in a corrected alpha of p < 0.05. Unthresholded statistical maps for the condition and time main effects, as well as the condition by time interaction can be found here: https://neurovault.org/collections/NLPFUQRB/.

## Results

### Adherence

We monitored the amount of time that the participants used the online program. The mean cumulative time dedicated to the intervention by our participants was 4885 minutes, with a standard deviation of 2470 minutes, while the average time spent per day was 66.5 minutes, accompanied by a standard deviation of 25.1 minutes.

### Behavioral Results

An ANOVA with time (pre and post-language learning) and condition (congruent and incongruent) as repeated measures with the accuracy data identified a significant main effect for condition, *F(1,40) = 25.373, p < 0.001*, characterized by increased accuracy on congruent trials relative to incongruent trials. There was also a significant main effect for time, *F(1,40) = 5.219, p = 0.028*, characterized by more accurate performance after language learning relative to before. We also found a significant condition by time interaction, *F(1,40) = 8.203, p = 0.007*, characterized by a greater difference in accuracy between congruent and incongruent trials before, *t = 5.756, p < 0.001*, and less difference in accuracy between congruent and incongruent trials after language learning, *t = 2.37, p = 0.061*, (Figure 1A).

**Figure 1.**
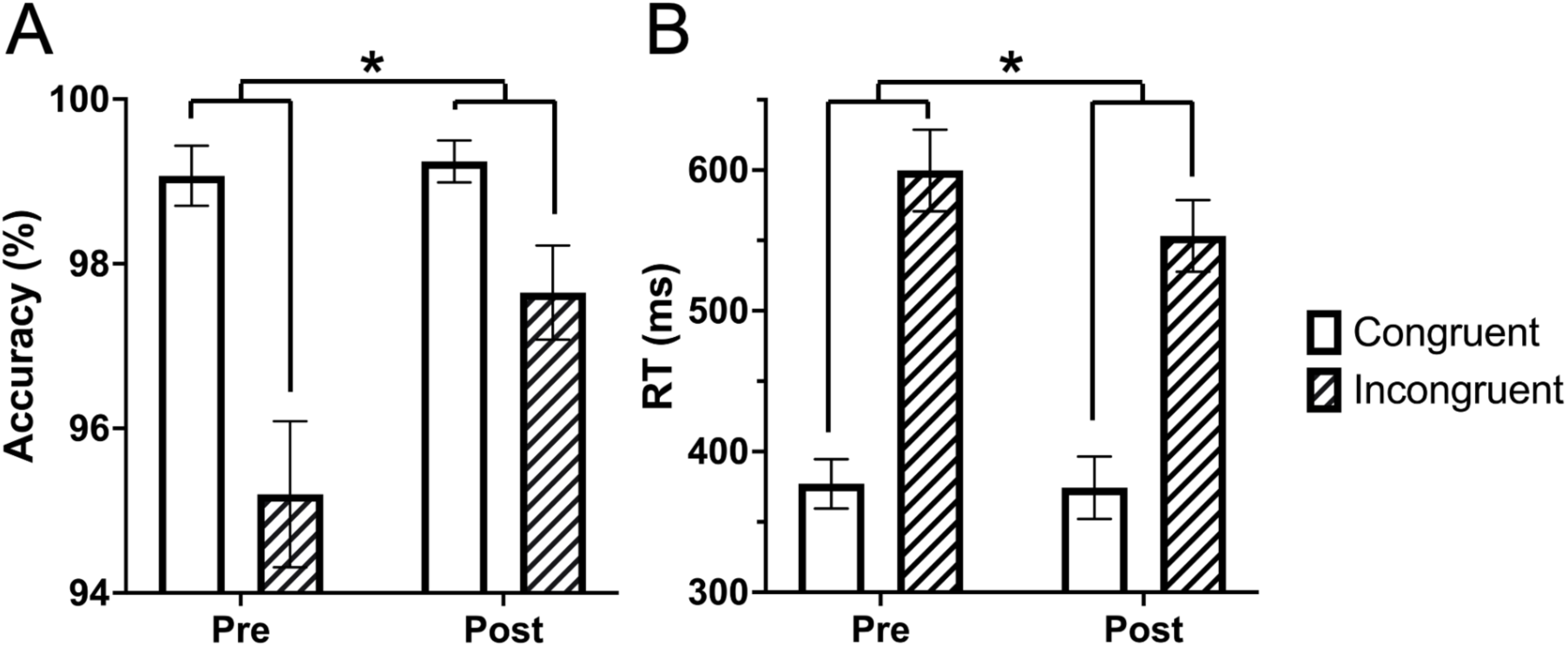
Performance on the Stroop task increases from pre to post language learning. (A) The difference between accuracy on congruent and incongruent trials decreases following language learning. (B) The difference between reaction time on congruent and incongruent trials decreases following language learning. Asterisks indicate interaction p-value < 0.05.

An ANOVA with time (pre and post-language learning) and condition (congruent and incongruent) as repeated measures with the reaction time data identified a significant main effect for condition, *F (1,40) = 218.813, p < 0.001*, characterized by faster reaction time on congruent trials relative to incongruent trials. We also found a significant condition by time interaction, *F (1,40) = 10.010, p = 0.003*, characterized by a difference in reaction time between incongruent trials pre-language learning and post-language learning with faster reaction time post-language learning, *t = 3.213, p = 0.004*, and no difference in reaction time between pre- and post-language learning on congruent trials, *t = 0.192, p = 0.849*. There was not a significant main effect for time, *F (1,40) = 3.752, p = 0.06*, (Figure 1B).

### fMRI Results

#### Stroop Main Effect

Voxel-wise analysis of the fMRI data was conducted using 3dMVM in AFNI. The multivariate modeling approach identified a main effect for condition (incongruent vs. congruent trials). All significant clusters were characterized by greater responses on incongruent trials relative to congruent trials. We identified significant differences between incongruent and congruent trials in portions of the lateral prefrontal cortex, parietal cortex, and the anterior cingulate, consistent with previous studies using the Stroop task (Huang et al., 2020; Mill et al., 2020; Song & Hakoda, 2015) (Figure 2, and Table 1).

**Figure 2.**
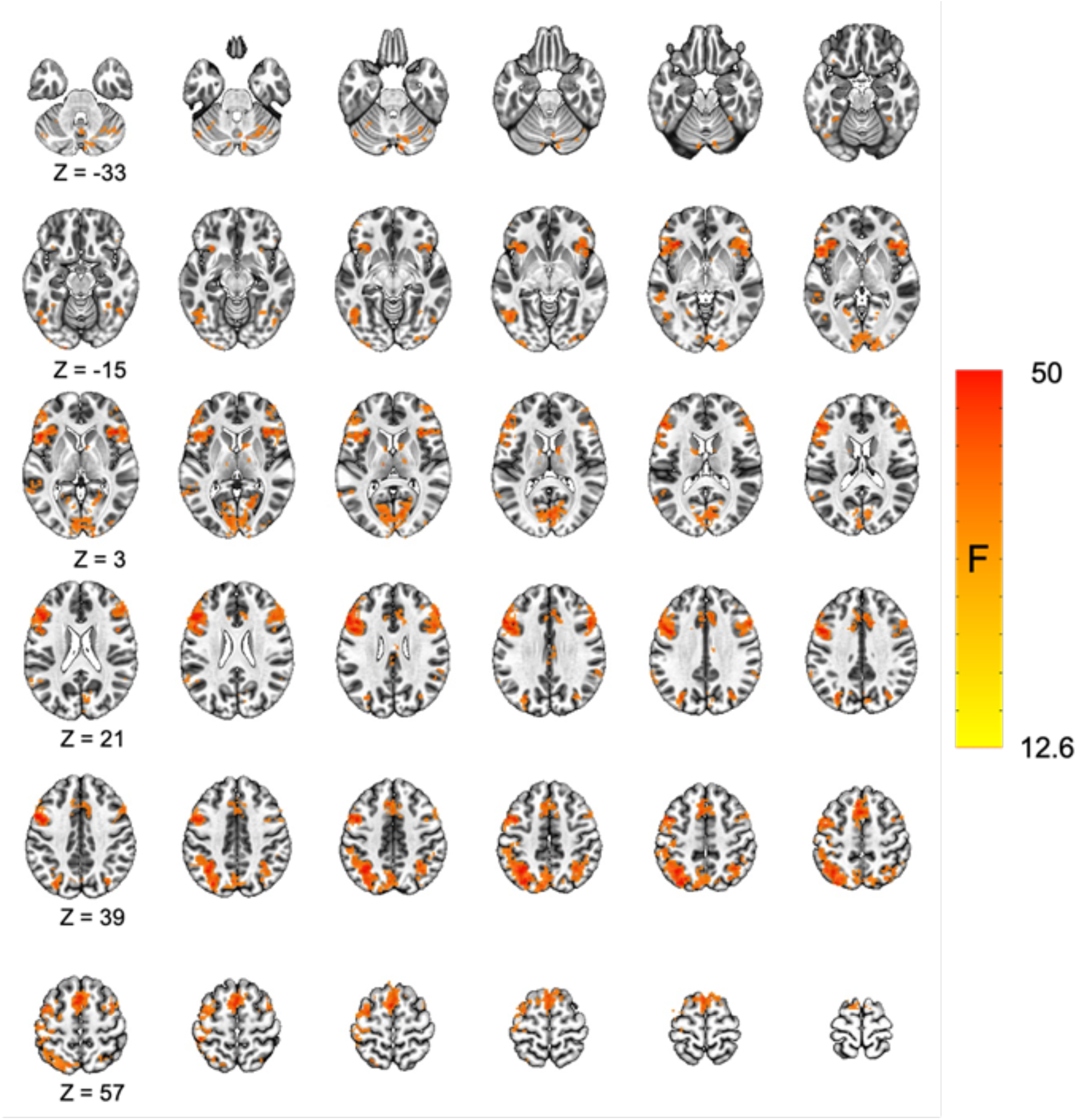
Increased activation for incongruent relative to congruent trials in the Stroop task. The main effect for condition (incongruent vs. congruent) is displayed. All significant results were characterized by greater activation for the incongruent relative to congruent condition. Cluster corrected p-value < 0.05.

**Table 1.**
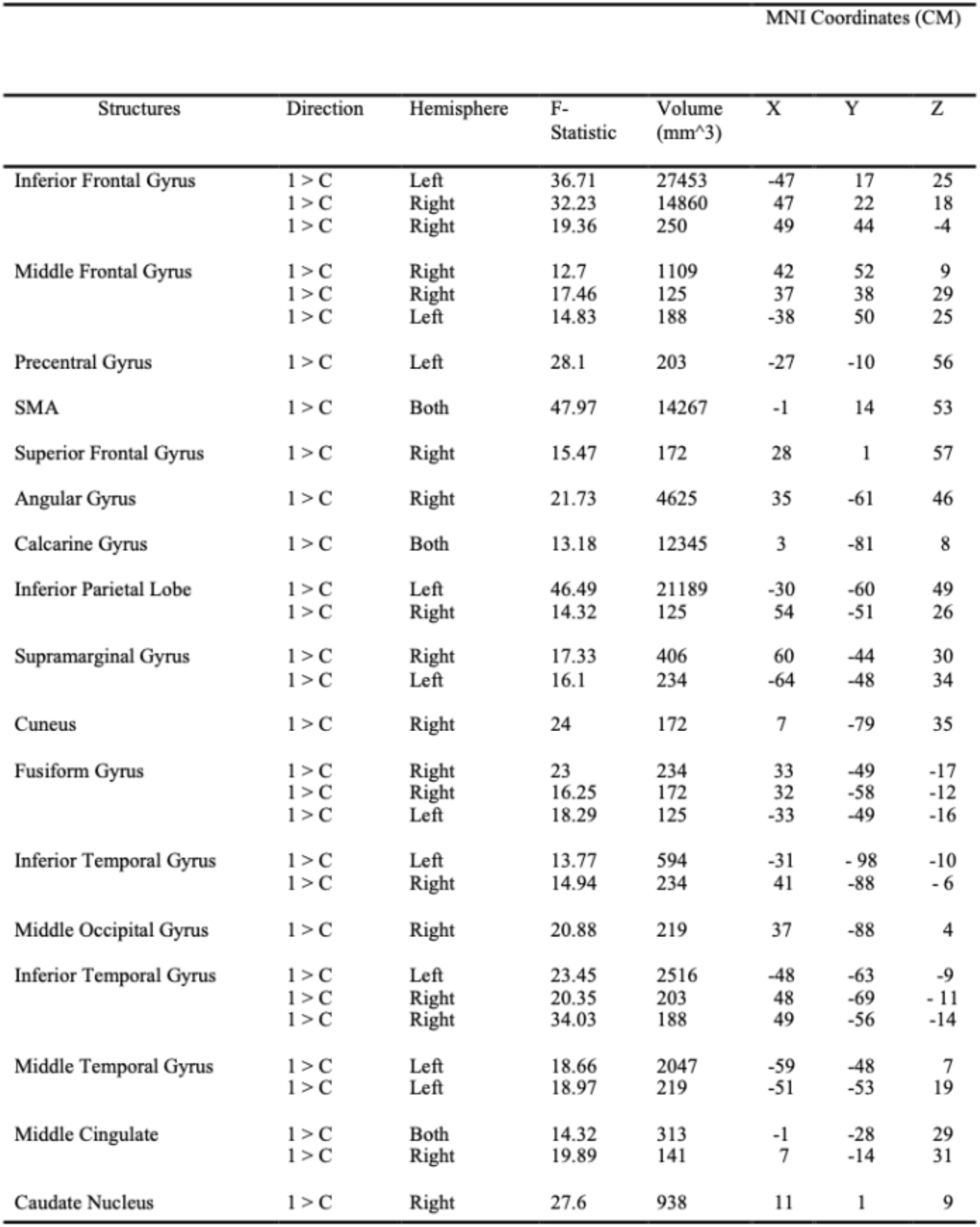

| MNI Coordinates (CM) |  |  |  |  |  |  |  |
| --- | --- | --- | --- | --- | --- | --- | --- |
| Structures | Direction | Hemisphere | F-Statistic | Volume (mm<sup>3</sup>) | X | Y | Z |
| Inferior Frontal Gyrus | 1 > C | Left | 36.71 | 27453 | -47 | 17 | 25 |
|  | 1 > C | Right | 32.23 | 14860 | 47 | 22 | 18 |
|  | 1 > C | Right | 19.36 | 250 | 49 | 44 | -4 |
| Middle Frontal Gyrus | 1 > C | Right | 12.7 | 1109 | 42 | 52 | 9 |
|  | 1 > C | Right | 17.46 | 125 | 37 | 38 | 29 |
|  | 1 > C | Left | 14.83 | 188 | -38 | 50 | 25 |
| Precentral Gyrus | 1 > C | Left | 28.1 | 203 | -27 | -10 | 56 |
| SMA | 1 > C | Both | 47.97 | 14267 | -1 | 14 | 53 |
| Superior Frontal Gyrus | 1 > C | Right | 15.47 | 172 | 28 | 1 | 57 |
| Angular Gyrus | 1 > C | Right | 21.73 | 4625 | 35 | -61 | 46 |
| Calcarine Gyrus | 1 > C | Both | 13.18 | 12345 | 3 | -81 | 8 |
| Inferior Parietal Lobe | 1 > C | Left | 46.49 | 21189 | -30 | -60 | 49 |
|  | 1 > C | Right | 14.32 | 125 | 54 | -51 | 26 |
| Supramarginal Gyrus | 1 > C | Right | 17.33 | 406 | 60 | -44 | 30 |
|  | 1 > C | Left | 16.1 | 234 | -64 | -48 | 34 |
| Cuneus | 1 > C | Right | 24 | 172 | 7 | -79 | 35 |
| Fusiform Gyrus | 1 > C | Right | 23 | 234 | 33 | -49 | -17 |
|  | 1 > C | Right | 16.25 | 172 | 32 | -58 | -12 |
|  | 1 > C | Left | 18.29 | 125 | -33 | -49 | -16 |
| Inferior Temporal Gyrus | 1 > C | Left | 13.77 | 594 | -31 | -98 | -10 |
|  | 1 > C | Right | 14.94 | 234 | 41 | -88 | -6 |
| Middle Occipital Gyrus | 1 > C | Right | 20.88 | 219 | 37 | -88 | 4 |
| Inferior Temporal Gyrus | 1 > C | Left | 23.45 | 2516 | -48 | -63 | -9 |
|  | 1 > C | Right | 20.35 | 203 | 48 | -69 | -11 |
|  | 1 > C | Right | 34.03 | 188 | 49 | -56 | -14 |
| Middle Temporal Gyrus | 1 > C | Left | 18.66 | 2047 | -59 | -48 | 7 |
|  | 1 > C | Left | 18.97 | 219 | -51 | -53 | 19 |
| Middle Cingulate | 1 > C | Both | 14.32 | 313 | -1 | -28 | 29 |
|  | 1 > C | Right | 19.89 | 141 | 7 | -14 | 31 |
| Caudate Nucleus | 1 > C | Right | 27.6 | 938 | 11 | 1 | 9 |

#### Time Main Effect and Condition by Time Interaction

We did not observe any significant clusters for the main effect of time (pre vs. post-language learning), and we did not find any significant clusters for the condition (incongruent and congruent) by time (pre and post-language learning) interaction.

#### Time Spent with the Language Learning Program is Related to Changes in Activity During Stroop Task

Next, we examined the possibility of whether the amount of time each participant spent using the language learning program had an effect on the changes in task-evoked activity from baseline to after the language learning intervention. First, we calculated a metric of change in activation over time. We calculated the difference score between incongruent and congruent conditions for each time point. Then we subtracted the pre- difference score from the post-language learning difference score ((post incongruent - post congruent) - (pre incongruent - pre congruent)). The result is a measure where higher values reflect a greater difference between incongruent and congruent post-language learning relative to pre. This measure was calculated for four regions (right and left lateral prefrontal cortex and right and left parietal cortex, see Figure 3A) that showed a main effect for task condition, and which have been previously implicated in performance during the Stroop task. Finally, we used Spearman rank correlation to examine the relationship between changes in activation from pre to post-language learning and the total amount of time each participant spent using the language learning program. The change in activation was positively correlated with time spent on the language learning program, *rho = 0.527, p < 0.001,* (Figure 3B), in the left prefrontal cortex and the left parietal cortex, *rho = 0.397, p = 0.01*, (Figure 3C). There was also a positive relationship between changes in activity and time spent on the language learning program in the right lateral prefrontal cortex, *rho = 0.454, p = 0.003*, (Figure 3D) and the right parietal cortex, *rho = 0.367, p = 0.018*, (Figure 3E).

**Figure 3.**
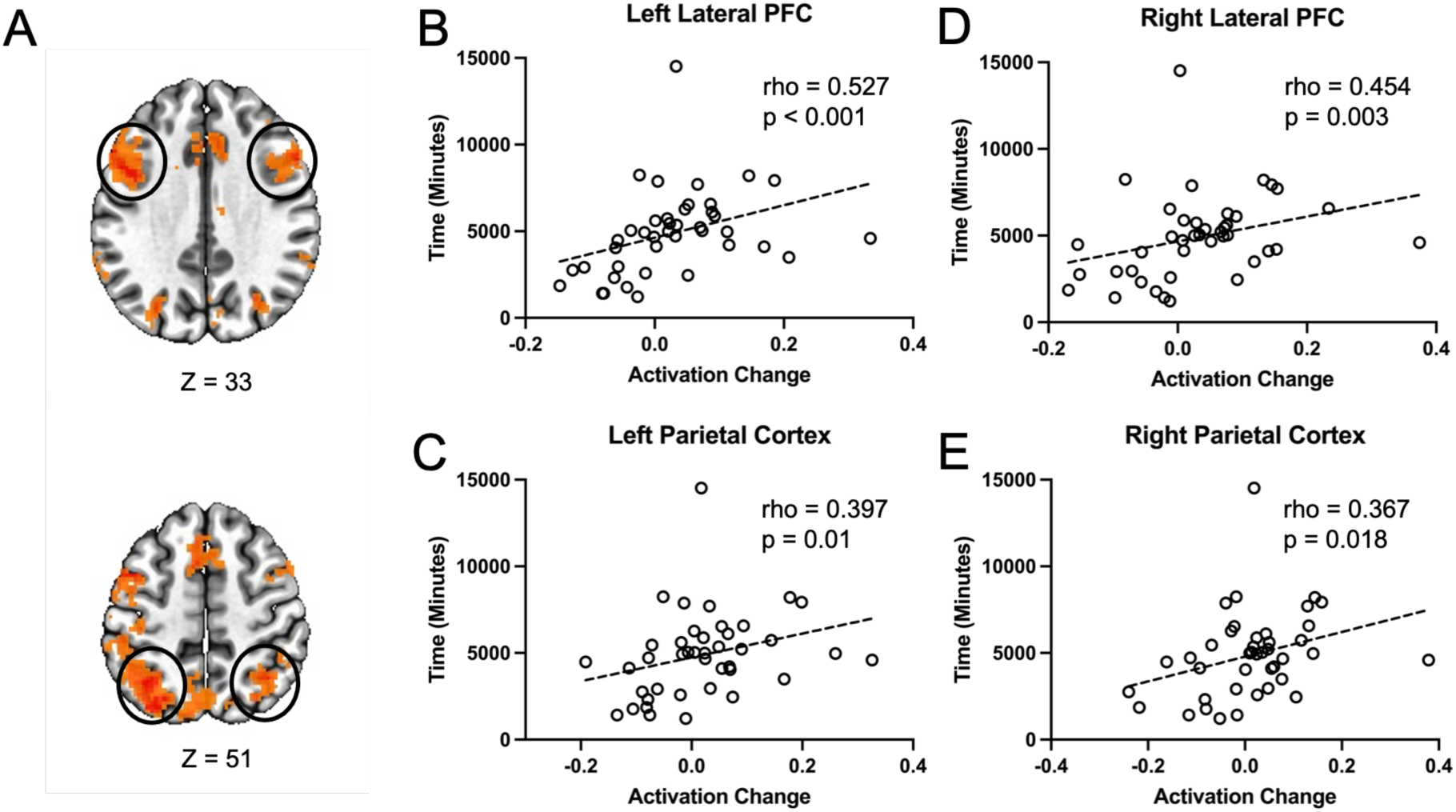
Changes in Stroop activation are related to the amount of time spent on the language learning program. (A). Changes in Stroop task activation were extracted from four clusters identified in the main effect for condition. These clusters were selected based on their involvement in the Stroop task in previous studies as well as from the main effect of condition in the current study. Correlation between the change in Stroop activity and the time participants spent with the language learning program in the (B) Left lateral prefrontal cortex, (C) Left parietal cortex, (D) Right lateral prefrontal cortex, and (E) Right parietal cortex.

We observed that changes in Stroop activation were related to the amount of time people spent on the language learning program. Next we examined whether time spent on the language learning program was related to the change in performance on the task. Based on the fMRI results, we hypothesized that increases in behavioral performance would be positively correlated with the amount of time spent on the language learning program. We calculated a behavioral change score for both accuracy and reaction time. This behavioral change score followed the same logic as that used for the fMRI activation change score. However, as improvements in performance over time result in more similar accuracy and reaction time for congruent and incongruent times (see Figure 1), we reversed the subtraction operations so that more positive scores could be interpreted as increases in performance from pre to post intervention for both accuracy and reaction time. Therefore the formula for accuracy was ((pre congruent - pre incongruent) - (post congruent - post incongruent)). Additionally, there was greater between-participant variability in reaction times, so to account for this variability we calculated the change score as a proportion of the difference between incongruent and congruent trials pre-intervention. Thus, the formula for reaction time was ((pre incongruent - pre congruent) - (post incongruent - post congruent)) / (pre incongruent - pre congruent)). Next we correlated the behavioral change scores with the amount of time spent on the language learning program. As our hypothesis was that this relationship would be positive, we used a one-tailed test. We did not observe a significant relationship between time spent on the language learning program and improvements in either accuracy or reaction time on the Stroop task, *smallest one-tailed p = 0.199*.

While we did not observe a significant linear relationship between time spent on the language learning program and improvements in Stroop performance, we split participants into two groups, those who spent more time with the language learning program and those who spent less time. Participants were placed in these groups if they spent more or less than the mean amount of time with the language learning program (4885 minutes or ∼81.5 hours). Then we evaluated whether or not these groups were characterized by different levels of improvement on Stroop task performance (as described above). We found that the group who spent more time with the language learning program showed greater increases in Stroop accuracy from pre to post-intervention, *t(39) = 1.846, one-tailed p = 0.036*. We did not observe any differences in reaction time performance between the groups who spent more or less time with the language learning program, *t(39) = -1.036, one-tailed p = 0.153*. These data suggest that time spent with the language learning program was related to greater accuracy gains from pre to post-intervention on the Stroop. While the effect on reaction time was not significant, it was in the opposite direction as the accuracy effect which may be suggestive of a speed/accuracy trade off.

## Discussion

We hypothesized that learning a new language could serve as a beneficial intervention to boost cognitive performance for older adults. By leveraging the cognitive complexity inherent in acquiring a new language, we hypothesized that L2 learning would lead to specific cognitive domains such as executive functions, such as enhanced cognitive control, and quicker information processing.

Behavioral results showed the Stroop effects and an improved performance post intervention as compared to pre intervention. Specifically, pre-post intervention comparisons reflected that the performance in Stroop task was more accurate post intervention as compared to pre intervention and the accuracy of the incongruent trials, which require better cognitive control, were improved and closer to the easier congruent trials which are less demanding in terms of conflict management. Response times showed a similar effect to accuracy suggesting language learning influenced processing speed as well. Together, these data support the idea that Stroop performance improves following language learning.

The functional neuroimaging results support the behavioral results. The activation maps regarding contrasts between incongruent and congruent trials are consistent with previous studies using the Stroop task (Huang et al., 2020; Mill et al., 2020; Song & Hakoda, 2015), with greater activations for the incongruent trials reflecting higher cognitive demand required for processing incongruent trials. The significant activations observed were in parts of the lateral prefrontal cortex, parietal cortex, and the anterior cingulate, all of which are involved in executive function, inhibitory control, conflict management and working memory (Ardila, 2019; De Pisapia et al., 2006; Friedman & Robbins, 2022), cognitive domains required for processing the Stroop task (Heidlmayr et al., 2020; Ye & Zhou, 2009).

While we did not observe any significant effect of time (pre vs. post-language learning), or for the interaction (condition by time), we did observe a relationship between changes in the brain activity during Stroop task and the amount of time participants spent on language learning.

Specifically, we found a positive relationship between changes in brain activity and time spent on the language learning program in the right lateral prefrontal cortex, the left prefrontal cortex and the left parietal cortex, as well as the right lateral prefrontal cortex. This suggests that the time spent on learning a new language can modulate activation related to the Stroop task in regions included in the frontoparietal network (FPN).

The FPN and cingulate play major roles in the organization of executive function, including working memory, conflict management, inhibitory control, and planning (Friedman & Robbins, 2022; Menon & D’Esposito, 2022). The FPN, with its flexible hubs, specially in the right hemisphere, is instrumental in modulating activity across various distributed systems, such as visual, limbic, and motor networks, to align with goals of task in hand, directly supporting working memory and task-switching capabilities (Cole et al., 2014). On the other hand, the cingulo-opercular network (CON) is crucial for maintaining task sets, enabling sustained cognitive performance across different tasks (Hausman et al., 2022). The CON functions through connections with other hub regions, including the anterior cingulate and anterior insula, which are associated with better performance in working memory, inhibition, and set-shifting tasks (Bush & Shin, 2006; Hausman et al., 2022). Together, these networks form part of a superordinate cognitive control system that includes the dorsolateral prefrontal, anterior cingulate, and parietal cortices, underpinning a broad array of executive functions (Niendam et al., 2012). Patterns of activity in regions of these networks can contain information about diverse task rules across multiple domains illustrating how they may function to orchestrate goal-directed behavior (Ito et al., 2017; Schultz et al., 2022). This superordinate network can describe how the executive function system with a common infrastructure can cater to the requirements of a network-specific and domain-specific demands of a particular task.

Both dorsolateral prefrontal and parietal cortex have been consistently reported as key brain areas associated with Stroop task performance that are involved in cognitive control and interference control (Huang et al., 2020). Dorsolateral prefrontal and parietal cortex are consistently implicated in Alzheimer’s disease and mild cognitive impairment (Giovacchini et al., 2019; Morbelli et al., 2013), and as critical brain areas related to cognitive reserve (Dodich et al., 2018; Giovacchini et al., 2019; Ye & Zhou, 2009). Interestingly, these brain areas have been reported as important brain areas that contribute to the cognitive advantage in lifelong bilinguals as compared to monolingual peers. The current results suggest that learning a new language in older adults for four months can improve Stroop performance. These results may also suggest that learning a new language in older adults can contribute to improving cognitive reserve and may contribute to postponing or slowing cognitive decline.

These results are consistent with studies that have explored a similar research question. Specifically, to date, there are four studies that have explored the cognitive effect of language learning in older adults (Bubbico et al., 2019; Meltzer et al., 2023; Nilsson et al., 2021; Wong et al., 2019). These studies use different measures and therefore the results are not easy to compare. Nevertheless, all four studies evidence some improvement post intervention (language learning) in a specific cognitive domain (Meltzer et al., 2023; Nilsson et al., 2021) or global cognition (Bubbico et al., 2019; Wong et al., 2019). Meltzer and colleagues (2023) reported a behavioral study in 76 older adults aged 65-75, similar to the results of our study, and found that Stroop accuracy improved after 16 weeks of either language learning with Duolingo or with an equivalent amount of non-language cognitive training. These results are consistent with what we observed in the current study. However, Meltzer and colleagues (2023) report that the non-language cognitive training condition resulted in improvements in reaction time on the Stroop while Duolingo did not. These results are in contrast to the current study where we did observe an effect of language learning on reaction time on the Stroop task. It is unclear whether potential differences in the dose, duration, intensity of the intervention, or an additional factor, may contribute to this difference.

Another study examined the neurocognitive effects of four-month second language learning program on the brain function of healthy elderly individuals (age 59-79) in a behavioral and neuroimaging study (Bubbico et al., 2019). They reported the behavioral impact as limited to global cognition. However, post-program, participants showed increased resting state functional connectivity in the right inferior frontal gyrus, right superior frontal gyrus, and left superior parietal lobule. Collectively, these brain areas are involved in numerous processes including: inhibitory control, decision making, attentional control, processing indirect language components such as emotional aspects, (Eriksen & Zacharov, 2016; Simonet, 2014), maintaining and manipulating information in working memory, as well as in directing attention (Manelis & Reder, 2014), and spatial attention and orientation (Heinen et al., 2017). In the current study, the contrast between incongruent and congruent trials on the Stroop task yielded activation of left lateral prefrontal cortex, and right lateral prefrontal cortex, as well as the left and the right parietal cortex. Although not a precise overlap, the activation map we observed is consistent with many of the regions showing changes in resting-state connectivity by Bubbico and colleagues (2019).

Altogether, in line with the previous reports (Bubbico et al., 2019; Meltzer et al., 2023), we provide evidence that even short-term language learning interventions can significantly reorganize and enhance neural network functions in older adults, contributing to better cognitive performance in specific executive function tasks such as the Stroop task. It is important, however, to note that while Stroop performance improved from pre to post-intervention and brain activity was modulated by the amount of time people spent with the language learning program, the dose dependent effects of time spent on the language learning program and improvements in Stroop performance were tenuous. Future studies should determine the optimal intensity, frequency, duration and intensity of language learning programs as an effective cognitive intervention in older adults.

We argue that language learning with sufficient dose, intensity and frequency can be considered as an effective cognitive intervention for promoting healthy aging. Based on our understanding of the cognitive reserve (Nelson et al., 2021; Stern, 2002, 2009, 2021; Stern et al., 2020) and the factors that can contribute to strengthening the cognitive reserve (Stern et al., 2020), this intervention may potentially counteract the cognitive decline associated with aging through the enhancement of brain plasticity and functional connectivity. The latter hypothesis remains to be tested.

## Conclusion

We bring evidence that language learning at older ages boosts cognitive control performance, as measured by improvements in the Stroop task. This enhancement is associated with functional neuroplasticity in cognitive control areas of the brain, indicating that acquiring a new language can actively influence function in these crucial regions. Specifically, these changes are observable improvements in tasks that require attentional control, such as the Stroop task, reflecting the transfer of cognitive gains from language learning to other cognitive domains. These results align with the theory of cognitive reserve, suggesting that intellectually stimulating activities like language learning can bolster the brain’s resilience to age-related decline. However, we also acknowledge that the extent of these benefits depends on the dose of language engagement, highlighting the importance of the amount and intensity of learning in realizing cognitive advantages. These results present a promising avenue for non-pharmaceutical intervention for aging related cognitive decline, offering an accessible, low cost and potentially enjoyable approach to maintaining and enhancing cognitive health in the aging population. As populations globally are experiencing increased longevity, understanding how to maintain cognitive health in later life is of paramount importance.

### Limitations and future research

Our study has a few limitations. First, our study’s design involved 41 participants, and is likely too small to allow for broad generalizations. Second, our study employed a pre-post intervention design without incorporating a control group or control condition, limiting our ability to isolate the intervention effect from other potential influencing factors such as practice effects. The challenges associated with conducting randomized trials on second language acquisition as a cognitive intervention in older adults, have limited research in this area. Future research approaches with bigger samples and designs that include control conditions are required. Third, this study potentially exhibits self-selection sample bias, as the personality types of the participants and demographic homogeneity could have influenced the composition of our sample. The participants were homogeneously white caucasian, which is reflective of the population characteristics of Nebraska. Future multi-site studies should include all races and different ethnicities. In addition, our sample consisted of older adults with moderate to high education levels (M=17.5, SD=2.94). Higher education level may have impacted the learning process or the cognitive demand required for learning a new language, and thus the results. Future studies should explore the effect of education level on the cognitive effect of language learning. Further, the ethnic minorities were excluded due to their proficiency in two or more languages. Future research should look at the cognitive effects of a third language in bilinguals of different ethnicities. Finally, our priority was to allow participants to select the language of their choice. This approach does not account for the effect of language distance. Future studies should explore the language distance effect. Thus, comprehensive research on cognitive effects of language learning is required to support our evidence for cognitive benefits of learning a new language in older adults. Further research can provide invaluable insights into cognitive aging, offering strategies not only for individual cognitive health but also for societal well-being.

## Acknowledgements

This study was funded by the University of Nebraska Collaboration Initiative Grant. Thank you to Rosetta Stone’s higher education division for providing complimentary access to their language learning curriculum and platform for the research participants, to our graduate students, Alexa Yunes-Koch, Annelie Personn, and Lauren Rezac, and to MRI technologist, Joanne Murray who all contributed to data acquisition, and to undergraduate student Noelle Abels for assistance in gathering and organizing references.

